# Pin1 Regulatory miRNAs as Novel Candidates for Alzheimer’s Disease Treatment

**DOI:** 10.1101/472985

**Authors:** Elyas Heidari, Elham Salehi Siavashani, Mohammad Rasooli, Zahra Amiri, Ali Sharifi-Zarchi, Koorosh Shahpasand

## Abstract

Alzheimer’s disease (AD) is the sixth leading cause of death in elderly people whose pathological hallmarks include senile plaques and neurofibrillary tangles (NFTs). The tangles are composed of hyperphosphorylated tau, which is a microtubule-associated protein and its hyperphosphorylation would result in its aggregation and neural cell death. Recently, it has been shown that phosphorylated tau at Thr231 exists in two distinct *cis & trans* conformations, whose conversion is being mediated by Pin1 isomerase and that the *cis*, but not the *trans*, is extremely neurotoxic and drives tau hyperphosphorylation. It has been demonstrated that Pin1 inhibition reflects *cis* pT231-tau accumulation in neurons but its overactivation is observed in cancer stem cells. Hence, a precise Pin1 regulation is required to keep cells in healthy conditions. As miRNAs play a crucial role in fine-tuning of the gene-expression level, we hypothesized that they might regulate the Pin1 dosage. Nonetheless, the possible regulatory roles of miRNAs in progression of AD by regulating PIN1 is not well studied. We aimed to identify potential miRNAs that down-regulate PIN1 in AD. This can uncover new regulatory mechanisms that result in AD. Thus, we performed a comprehensive study of miRNAs, capable in regulating Pin1, through whole-genome meta-analysis by integrating miRNA expression profiles of 846 different biological samples, along with a systematic literature review and data mining of multiple experimental and predicted miRNA-target databases. We created a list of 56 candidates, which was then short-listed to 10 miRNAs with vigorous experimental evidence. We examined the expression patterns of these miRNAs in the AD and healthy controls and integrated mRNA and miRNA expression profiles to study possible interactions between miRNAs and Pin1. Moreover, we performed an *in-silico* functional analysis by integrating data of knock-in and knock-down experiments of the candidate miRNAs, and highlighted miR296-5p, miR200b, miR200c, miR140-5p, and miR874 as strong candidate Pin1 regulators. These findings would have profound implications in developing novel therapeutic strategies for AD by blocking expression of highlighted miRNAs using antagomirs.

## I. INTRODUCTION

Alzheimer’s disease (AD) is a chronic, progressive, age-related neurodegenerative disorder which is the first cause of dementia worldwide. There are around 47 million AD patients globally which is 10–30% of the elderly population (Masters et al., 2015;Prince et al., 2016). Two main pathological hallmarks of AD are β-Amyloid Plaques and Neurofibrillary Tangles (NFTs) composed of Aβ and hyperphosphorylated tau respectively (Holtzman, Bales, & Paul, 2002;Laferla, 2002;Masters et al., 2015).

Tau is a phosphoprotein with around 85 phosphorylation sites and is essential for the assembly, structure and stability of microtubules (Brunden, Trojanowski, & Lee, 2009;Wang & Mandelkow, 2015). It is moderately phosphorylated under physiological conditions, but its hyperphosphorylation would result in pathogenicity and NFT formation. Despite extensive work, it was obscure which phosphorylation site is the most critical for driving the pathogenicity. Recently, we found that phosphorylated tau at Thr231 exists in the two distinct *cis* and *trans* conformation and that *cis* pT231-tau is extremely neurotoxic and drives the tauopathy, a process termed *cistauosis* (Kondo et al., n.d.).

Pathogenic *cis* pT231-tau is being normally converted to physiologic *trans* epitope by Peptidylprolyl Isomerase, NIMA-Interacting 1 enzyme (Pin1), which is an endothermic reaction (Nakamura et al., 2012). Pin1 functions like a double-edged sword. On the one hand, Pin1 down-regulation is shown in tauopathies including AD, Parkinson Disease, Traumatic Brain Injury (TBI) and brain stroke (Ballatore & Trojanowski, 2007;Mandelkow & Mandelkow, 2012) which in turn may result in *cistauosis*. On the other hand, Pin1 overactivation is associated with cancer stem cells (Zhou & Lu, 2016). Hence, a precise regulation of Pin1 is of crucial importance.

Most of the neuro-genetic studies have focused on the protein-coding genes and thus, possible roles of non-coding RNAs (ncRNAs) in this area remain highly obscure (Salta & Strooper, 2012). There are increasing reports about association of ncRNAs with memory, cognitive and behavioral processes (Santa-maria et al., 2015). miRNAs are a subset of endogenously initiated, small single-stranded non-coding RNAs regulating gene expression in post-transcriptional stage via either translational repression or mRNA degradation (Cai, Yu, Hu, & Yu, 2009). The regulatory role of miRNAs is claimed to be fine-tuning of the expression levels of the target genes, rather than switching them on or off (Lewis, Shih, Jones-rhoades, Bartel, & Burge, 2003). They are used as diagnostic biomarker in AD (Denk, Boelmans, Siegismund, Lassner, & Arlt, 2015;Kumar et al., 2013;Moon et al., 2016) and there are several reports about mediatory role of miRNAs in AD development (Hu et al., 2016;Rustighi et al., 2016;Yao, Guo, Reinhardt, Wulf, & Lieberman, 2015). However, potential function of miRNAs for regulating Pin1 in AD has not been scrutinized so far.

Herein we focus on the miRNAs which are known/hypothesised to bind Pin1, thereby regulating its expression level. We extract a set of candidate miRNAs from two online miRNA target prediction databases, namely, TargetScan7.1 and miRWalk2.0. Afterwards, we analyze a number of publicly available microRNA/RNA expression profiles and review the literature systematically to see if there is any evidence for each candidate regulation.

We believe our findings would have profound implication in AD therapy through Pin1 expression control, likely employing Antagomirs.

## II. MATERIALS AND METHODS

### • Pin1-targeting miRNAs

In order to find potential interactions between miRNAs and Pin1, we performed a systematic search over two miRNA-mRNA interaction databases including TargetScanHuman Release 7.1 (Agarwal, Bell, Nam, & Bartel, 2015) and miRWalk2.0 (Dweep, Sticht, Pandey, & Gretz, 2011) (Table 1). miRWalk2.0 predicts the binding region of each miRNAs to target genes by combining the predicted binding sites collected from 12 famous miRNA-target prediction software namely, DIANA-microTv4.0, DIANA-microT-CDS, miRanda-rel2010, mirBridge, miRDB4.0, miRmap, miRNAMap, doRiNA, PicTar2, PITA, RNA22v2, RNAhybrid2.1, and Targetscan6.2 based on biological analysis of promoter, cds, 5’-, and 3’UTR regions. Additionally, miRWalk2.0 integrates verified results from previous experiments through information from well-known databases miRTarBase, PhenomiR, miR2Disease, and HMDD (Dweep et al., 2011).

**Table 1.**
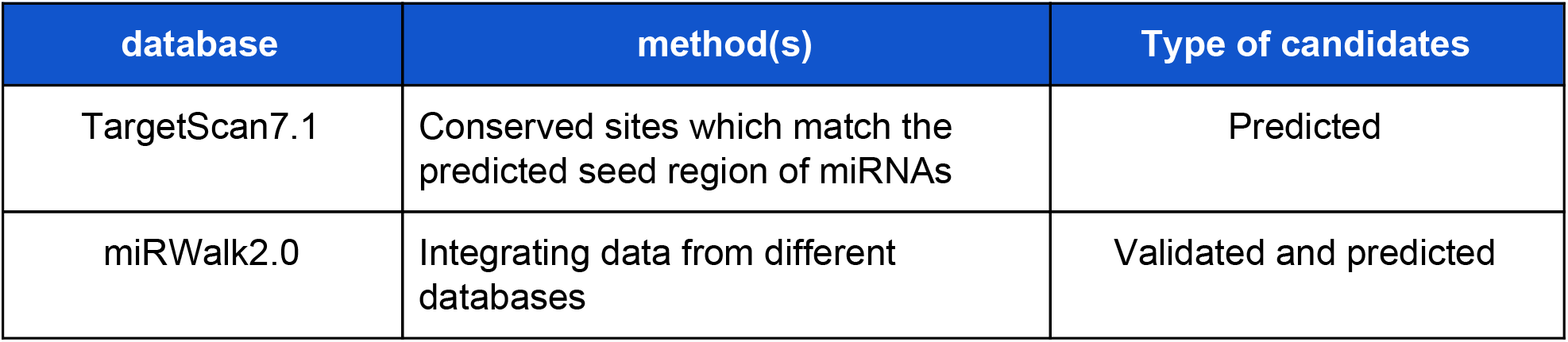
Utilized miRNA-mRNA interaction databases

| database | method(s) | Type of candidates |
| --- | --- | --- |
| TargetScan7.1 | Conserved sites which match the predicted seed region of miRNAs | Predicted |
| miRWalk2.0 | Integrating data from different databases | Validated and predicted |

TargetScan7.1 predicts miRNA-mRNA regulatory interactions through analysis of conserved 8mer, 7mer, and 6mer regions matching the seed region of miRNAs. Furthermore, the new version has been improved compared to the older ones (i.e. TargetScan6.2) to predict interaction efficacy, using 3’UTR and some other information (Agarwal et al., 2015).

### • Systematic review of extracted miRNAs

After extracting candidate miRNAs from the databases, we tried to find any evidence in the past literature for each of the interactions found in the previous step. According to the large amount of research done on Pin1 and work conducted on miRNAs, reviewing the literature manually could be exhaustive; hence, we developed an automated machinery so as to execute systematic review on the literature.

The R package RISmed was used to carry out systematic review on the abstracts available on PubMed. In this regard, we searched for nine terms “Pin1”, “Alzheimer’s”, “over-expression”, “induce”, “inhibit”, “transfect”, “knock-out”, “knock-in”, and “knock-down” along with official IDs of the extracted miRNAs from the aforementioned online databases (e.g. “hsa-mir-296-5p pin1”). Afterwards, the number of words between IDs and terms in a context was considered as a measurement of the importance and sharpness of a search result. Thereafter, if the distance between an ID and terms in a context of a search result was more than 15 words the result was excluded from analysis step.

Associated with the previous step, the online database Coremine containing natural language processing results on pertinent literature was used to achieve a similar goal. The keywords “Pin1” and “miRNA” were searched simultaneously in the database.

### • miRNA expression profiles in AD

A miRNA may be recognized to play a role in AD only if it is expressed differently at some level among AD and normal samples. In order to check that whether any of the candidate miRNAs express differentially in AD cases and normal controls and to what extent, we sought for miRNA expression profiles from different parts of the brain.

We collected miRNA expression data from 4 studies including 262 diagnosed AD-affected as well as 235 healthy controls (Cheng et al., 2014;Hu et al., 2016;Lau et al., 2013;Leidinger et al., 2013;Tan et al., 2014). The collected data were acquired from CSF, Hippocampus, plasma, and serum. Furthermore, we integrated the data of two additional experiments with 22 AD-affected samples and 28 controls from CSF and 4 AD-affected samples and 4 controls from parietal lobe (Denk et al., 2015;Nunez-iglesias, Liu, Morgan, Finch, & Zhou, 2010). Details about the experiments is presented in table 2.

**Table 2.** Included studies of miRNA-expression profiling in AD cases and normal controls.

| Reference | #Cases | #Controls | Specimen |
| --- | --- | --- | --- |
| Leidinger et al. (2013) | 48 | 22 | Plasma |
| Tan et al. (2014a) | 158 | 155 | Serum |
| Cheng et al. (2014) | 15 | 35 | Serum |
| Lau et al. (2013) | 41 | 23 | Hippocampus |
| Denk et al. (2015) | 22 | 28 | CSF |
| Zhou et al. (2010) | 4 | 4 | Parietal lobe |

After background-correction and normalization applied to raw data, we used limma package (R software) in order to calculate the differences in expression (Ritchie et al., 2015).

### • Integrated expression profiles of mRNAs and miRNAs

To test a regulatory interaction between the candidate miRNAs and Pin1, we collected both mRNA and miRNA expression profiles of a group of biological samples from the Gene Expression Omnibus database (GEO) (Table III) (Barrett et al., 2013). Then we computed correlations between the expression levels of Pin1 mRNA and the candidate miRNAs. We expected that a miRNA with inhibitory effect should be negatively correlated with the target gene.

**Table 3.** Summary information about miRNA-mRNA integrated databases

| ID | Specimen | #Samples |
| --- | --- | --- |
| GSE34608 | Pulmonary tuberculosis and sarcoidosis | 20 |
| GSE19783 | ER+/ER- breast cancer | 40 |
| GSE38226 | Liver fibrosis | 21 |
| GSE19536 | Breast cancer | 100 |
| GSE28260 | Renal cortex and medulla | 13 |
| GSE42095 | Differentiated embryonic stem cells | 23 |
| GSE26953 | Aortic valvular endothelial cells | 24 |
| GSE21032 | Prostate cancer | 83 |
| GSE32688 | Pancreatic cancer | 32 |
| GSE38974 | Chronic obstructive pulmonary disease | 25 |
| GSE21687 | Ependynoma primary tumors | 64 |
| GSE17498 | Multiple myeloma | 40 |
| GSE35602 | Colorectal cancer stromal tissue | 25 |

For this purpose, we developed a crawler in python and R programming languages, called miRawler. It finds proper GEO data-series containing expression profiles of target genes and candidate miRNAs, obtains both mRNA and miRNA expression profiles, calculates correlation matrix for each data series, and draws a heatmap of correlation matrix and a linear plot for each data-series (source code can be obtained from https://github.com/EliHei/Alzo/tree/master/miRawler). Table 3 summarizes 15 used GEO data-series.

### • Functional effect of candidate miRNAs on Pin1 expression

To experimentally check whether candidate miRNAs have functional effect on the expression of Pin1, we searched for keywords “over-expression”, “induce”, “inhibit”, “transfect”, “knockout”, “knockin”, and “knockdown” to find any mRNA expression profiles of cells treated with forced expression or inhibition of candidate miRNAs.

Detailed information about mRNA and miRNA expression profiles is reported in table 5. Differential expression of Pin1 between different conditions of the treated cells was calculated using limma package after background-correction and normalization of the raw data.

**Table 4.**
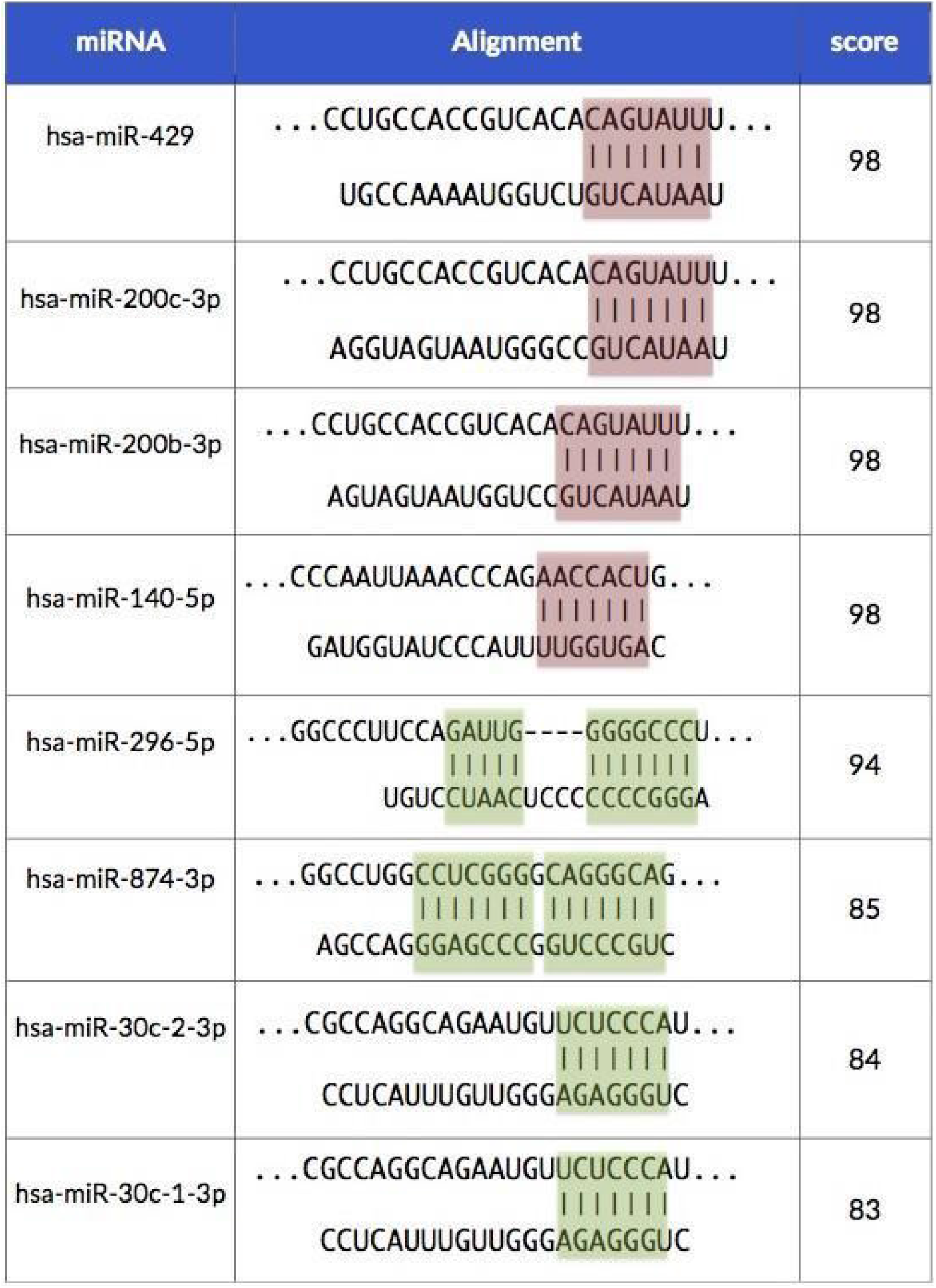
Binding sites of miRNA- Pin1 interactions based on TargetScanHuman. The upper sequence is a part of the Pin1 3’UTR and the lower sequence is the targeting miRNA. The score column indicates the *context*++ *score percentile* which is an index defined by TargetScanHuman in order to predict targeting efficiency based on 14 factors [22]. Red boxes illustrate the strongly conserved binding regions.

**Table 5.** Differential Expression of Pin1 in targeted cells vs normal controls.

| ID | Summary | p.value | logFC (control - case) |
| --- | --- | --- | --- |
| GSE24289 | dox-induced miR-200c/141 | 0.52 | -0.09400886 |
| GSE36565 | transfected with Pre-miR-30 and KD-miR-30 family. | 0.024 | -0.24850877 |
| GSE47937 | A virus infected in the presence of exogenous miR-147b, miR-190b, miR-199a, miR-512-5p, and miR-874 with 3 biological replicates for each group. | 0.876 | 2.32E-02 |
| GSE50422 | miR30-treated | 0.0374 | 0.19149452 |
| GSE56973 | miR-429 treated | 0.399 | 0.19247517 |
| GSE58860 | transfected Caco2 cells by MIR429 | 0.09842567 | 0.22901243 |
| GSE59883 | Colorectal cancer cell lines were transduced with lentiviral antisense miRNA inhibitors (miRZIPs) targeting miR-429 and miR-200b. | 0.96 | -0.0072575 |
| GSE62230 | Expression of miR-200c in claudin-low breast cancer | 0.000228 | 0.567 |
| GSE75800 | miR-199 as 23mer miRNA mimics induced. | 0.703 | -0.03238685 |
| GSE85629 | miR-199a expression was enforced in HMLE cells. | 0.47311 | -0.20931181 |

## III. RESULTS

### • Finding candidate miRNAs

We have extracted total 56 miRNAs through two online databases (supplementary table 1). Among which 10 miRNAs have strong clinical representations: miR199a-5p, miR199b-5p, miR200b-3p, miR200c-3p, miR-874-3p, miR429, miR296-5p, miR30c-1-3p, miR30c-2-3p, miR140-5p. Notably miR296-5p and miR429 have the utmost evidences in literature (Table VII). There have been two results for miR200b and miR200c within Coremine framework, which we have noticed in our systematic review as well.

**Table 6 (A&B).** Details of differential expression of miRNAs between affected and control. Log fold change (LogFC) has been computed such that AD - HC.

|  | Lau et al. (hippocampus) |  | Lau et al. (prefrontal cortex 1) |  | Lau et al. (prefrontal cortex 2) |  | Leidinger et al. |  | Cheng et al. |  |
| --- | --- | --- | --- | --- | --- | --- | --- | --- | --- | --- |
| ID | logFC | p.value | logFC | p.value | logFC | p.value | logFC | p.value | logFC | p.value |
| hsa-miR-199a-5p | 0.03886 | 0.774767 | 1.442377 | 0.0001062 | 0.2038601 | 0.0887414 | 0.451426 | 0.0462335 | -0.071126 | 0.6399515 |
| hsa-miR-199b-5p | 0.05267 | 0.602685 | 0.7132753 | 0.0203652 | 0.1037857 | 0.2774222 | NA | NA | 0.4982414 | 0.0382858 |
| hsa-miR-200b-3p | 0.30518 | 0.091955 | 0.0963898 | 0.7323166 | 0.0803924 | 0.3078379 | - | - | 0.8439471 | 0.1155711 |
| hsa-miR-200c-3p | 0.26118 | 0.261518 | 0.497646 | 0.0593873 |  |  | - | - | 0.1440922 | 0.5613724 |
| hsa-miR-2392 | - | - | - | - | - | - | - | - | - | - |
| hsa-miR-429 | 0.01606 | 0.860847 | 1.298597 | 0.081249 | -0.360256 | 0.0503146 | - | - | -0.164603 | 0.5185476 |
| hsa-miR-296-5p | -0.0950 | 0.244594 | 0.147622 | 0.6193779 | 0.2549204 | 0.0019657 | 0.377475 | 0.0085751 | -0.2678925 | 0.5430875 |
| hsa-miR-30c-1-3p | 0.17847 | 0.055394 | 0.3020222 | 0.0853717 | 0.2846869 | 0.0604712 | NA | NA | 0.4085226 | 0.0219536 |
| hsa-miR-30c-2-3p |  |  | -0.382140 | 0.0488181 |  |  | - | - | -0.192716 | 0.0879674 |

|  | Zhou et al. |  | Denk et al. |  | Tan et al. |  |
| --- | --- | --- | --- | --- | --- | --- |
| ID | logFC | p.value | logFC | p.value | logFC | p.value |
| hsa-miR-199a-5p | -1.169640 | 0.1067763 | 0.0433320 | 0.1586340 | 0.6298396 | 0.0245221 |
| hsa-miR-199b-5p | 2.0475475 | 0.0651494 | - | - | 0.6665741 | 1.41e-08 |
| hsa-miR-200b-3p | 0.3485621 | 0.4330939 | NA | > 0.99 | - | - |
| hsa-miR-200c-3p | 1.2519146 | 0.0615539 | NA | > 0.99 | - | - |
| hsa-miR-2392 | - | - | NA | > 0.99 | - | - |
| hsa-miR-429 | -0.484403 | 0.61213 | NA | > 0.99 | 7.0802312 | 0.0001281 |
| hsa-miR-296-5p | 1.27599 | 0.0105845 | -0.042712 | 0.2060432 | - | - |
| hsa-miR-30c-1-3p | -0.155638 | 0.5306259 | NA | > 0.99 | -1.010063 | 0.129629 |
| hsa-miR-30c-2-3p |  |  | NA | > 0.99 | -1.186852 | 0.0006858 |

**Table 7.**
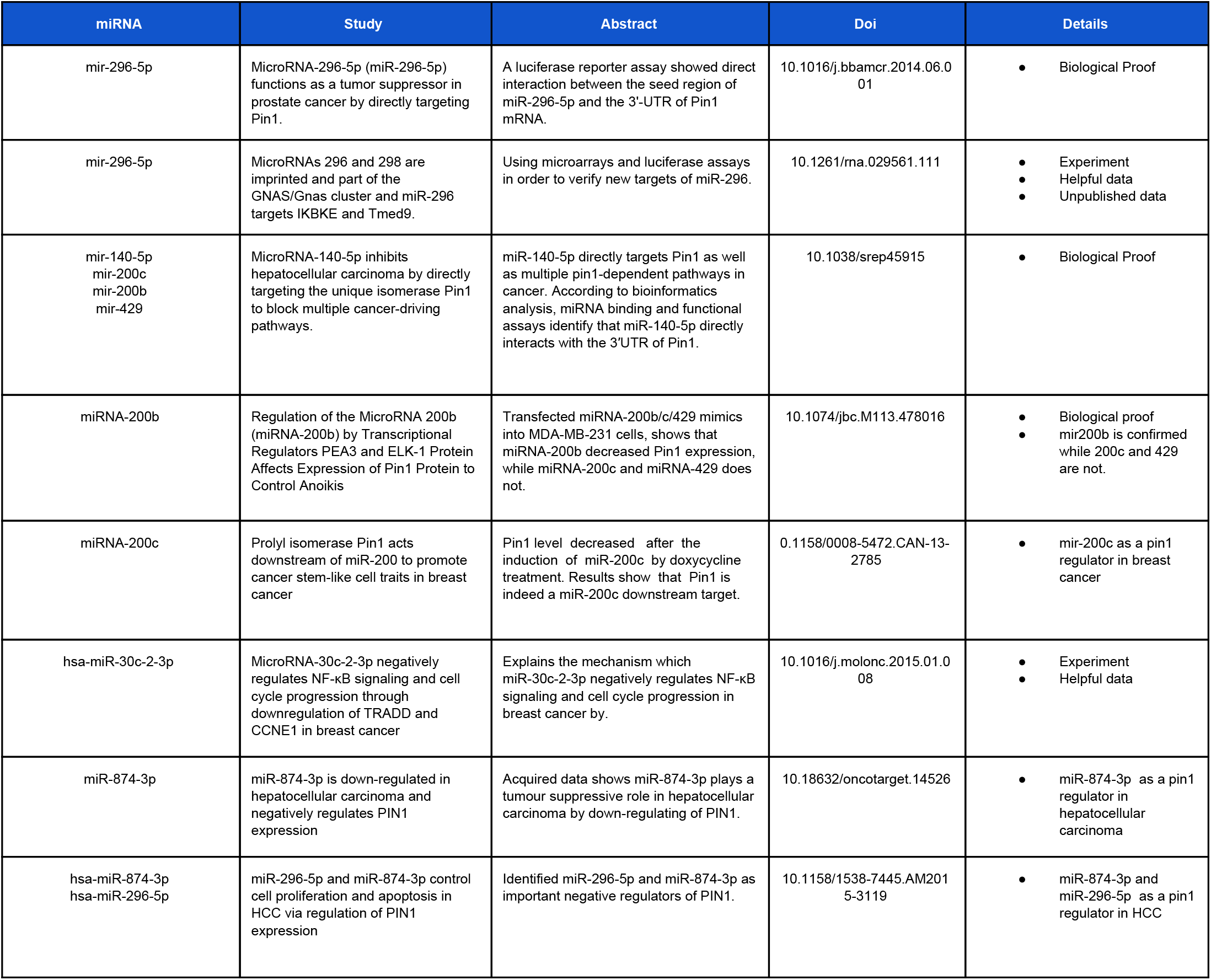
Remarkable findings in systematic review

Having eliminated the ones with no evidence, we examined the miRNAs sequences and Pin1 to find the inter-connections. Table 4 indicates the binding sites predicted by TargetScan7.1 program. The reported score is based on 14 features such as site type 3’UTR, minimum distance, seed paring stability, probability of conserved targeting, etc.

### • Candidate miRNAs expression patterns

We have initially calculated p-values and log fold changes of candidate miRNAs in preprocessed profiles of previous reports (Cheng *et al*., Zhou *et al*., and Lau *et al.*) (Ritchie et al., 2015). In addition, supplementary data of Leidinger *et al*., Tan et al., and Denk *et al*. were added to the table. We have found significantly different expression patterns of miR296-5p, miR199a-5p, miR199b-5p, and miR429 between healthy and AD brains; demonstrating the candidates mediatory role in AD.

### • Correlation between candidate miRNAs and Pin1

We showed a Pearson correlation between expression levels of candidate miRNAs and Pin1 (supplementary table 2). Expression profiles of GSE34608 and GSE28260 show significant negative correlation between candidate miRNA9s and Pin1. We demonstrated an inhibitory role of the candidate miRNAs in Pin1 expression (Fig. 2). Taking these together miR200b and miR296-5p are likely potent to Pin1 negative control.

**Figure 1.**
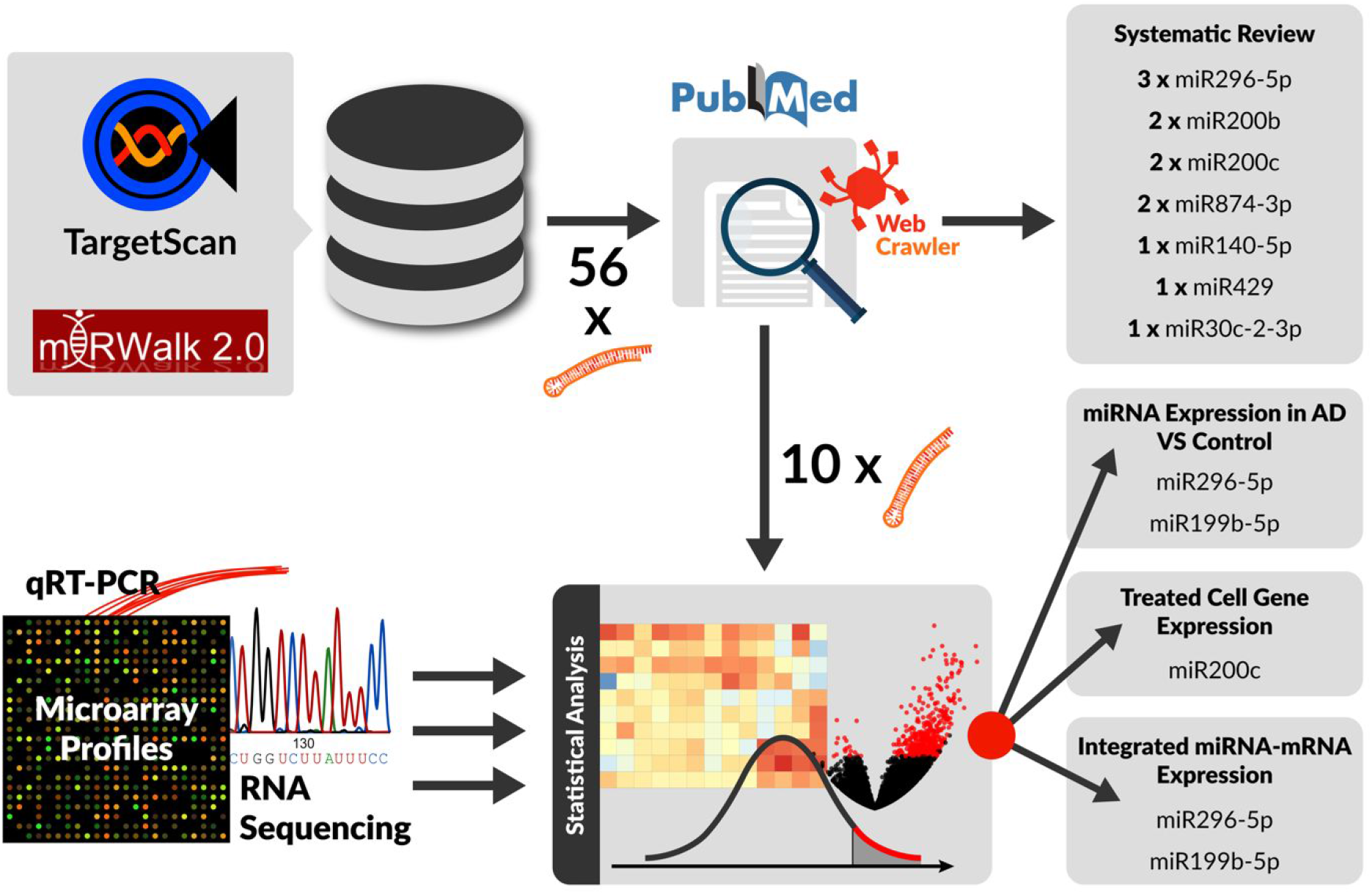
Overview of our pipeline. After extracting candidate miRNAs targeting Pin1 from TargetScan and miRWalk 2.0 databases, a systematic review on PubMed was carried out to find evidences on each candidate interaction, in which 10 candidates were shortlisted for further analyses. GEO datasets as well as a few other public datasets were used for statistical analyses. Our investigation, led to a list of miRNAs indicated in the figure as our final candidates to target Pin1.

**Figure 2.**
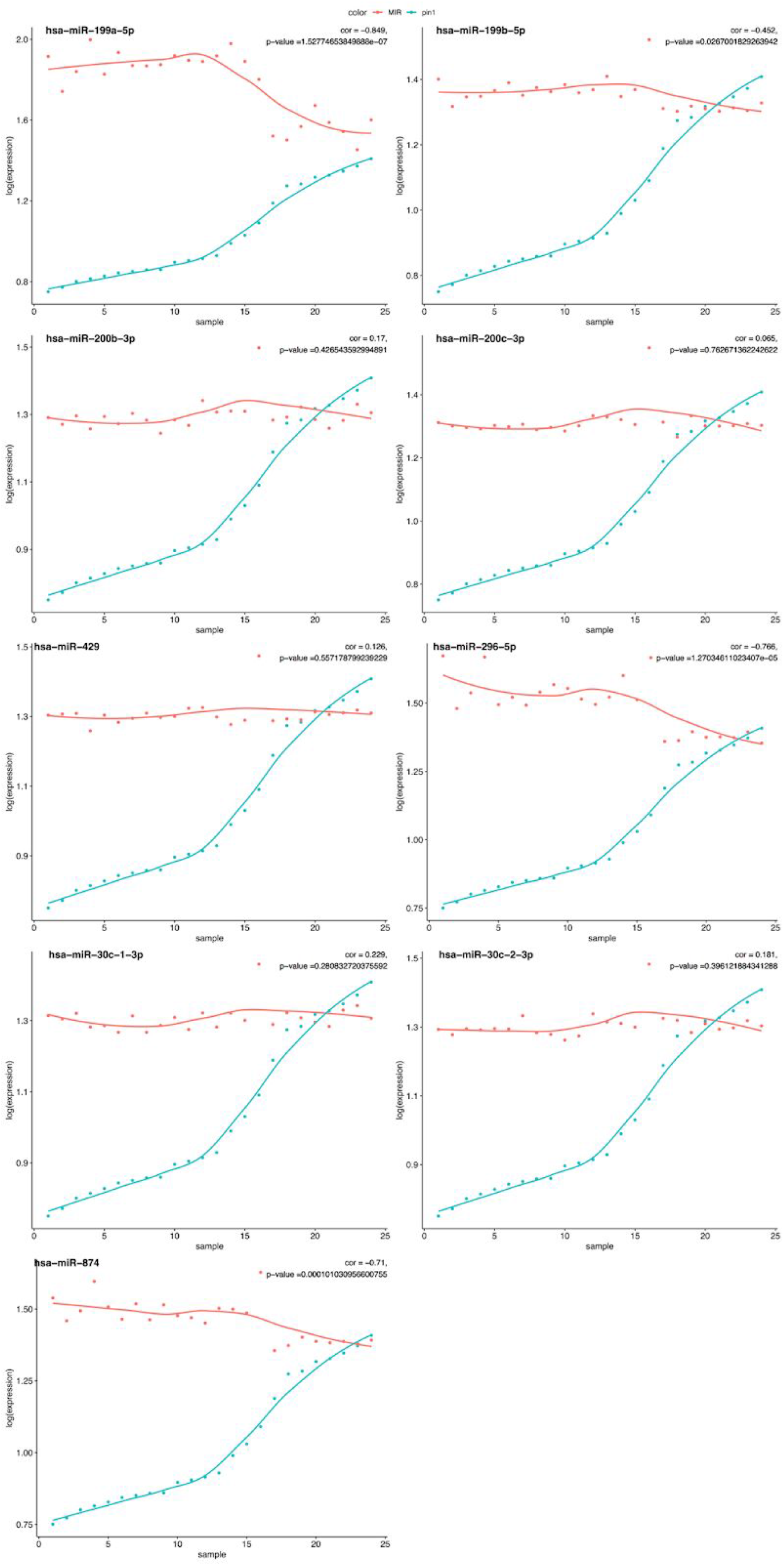
Negative expression correlation between PIN1 and the candidate miRNAs in GSE34608. Expression of candidate miRNAs as well as Pin1 gene in GSE34608. For each plot, x-axis represents different samples (sorted according to PIN1 expression level), and y-axis shows the mean-centered log2-scaled expression levels of PIN1 or a candidate miRNA (legend). Each of the blue and red points show the expression levels of a candidate miRNA and PIN1, respectively, in one sample. The blue and red curves depict the overall expression trend of the candidate miRNA and PIN1, respectively. Pearson correlation values between the expression of PIN1 and each candidate miRNA are shown above each panel.

### • Remarkable findings in systematic review

Sixteen discoveries from systematic review (apart from cell-treatment experiments) had valuable information about the problem in hand (table 3). We have found that mir-296-5p negatively regulate Pin1 expression in prostate cancer cells (Lee et al., 2014). Similarly, miR-296- 5p as well as miR-874-3p have been identified as important negative Pin1 regulators in hepatocellular carcinoma (HCC). Furthermore, Leong et al. findings introduced miR-874-3p as a tumor suppressor in hepatocellular carcinoma through Pin1 down-regulation (Leong, Cheng, Wong, & Ng, 2017). Yan et al. have recently demonstrated that mir-140-5p, which has not been reported as a predicted miRNA based on our selected online databases, inhibits HCC by Pin1 direct targeting but we couldn’t identify its effect within our selected online databases (Yan et al., 2017). miRNA-200b decreased Pin1 expression after transfecting miRNA-200b, miRNA-200c, and miRNA-429 mimics into MDA-MB-231 cells, while miRNA-200c and miRNA-429 did not (Zhang, Zhang, Gao, Wang, & Liu, 2013). Moreover, Luo et al. have shown that level of Pin1 levels is being decreased upon miR-200c induction by doxycycline (Yao et al., 2015).

## IV. DISCUSSION

Alzheimer Disease (AD) is an inclusive neurodegenerative disorder, which its economic burden has been around $226 billion in 2015. Despite of extensive considerations, an efficient therapeutic has been out of reach thus far. It has been recently shown that phosphorylated tau at Thr231 exists in the two distinct *cis* and *trans* conformations, whose conversion is being mediated by Pin1 isomerase and that *cis*, but not *trans*, is extremely neurotoxic and drives neurodegeneration upon tauopathies. Furthermore, they have demonstrated that Pin1 inhibition would reflect *cis* p-tau accumulation in neurons (Kondo et al., n.d.). Thus, it is of crucial importance to consider the factors regulating Pin1 expression and function.

miRNAs are being classified as non-coding RNAs and a class of regulatory factors mediating RNA silencing and post-transcriptional protein expression control. There are several reports claiming miRNAs participation in AD process. Moreover, various studies have demonstrated miRNAs as AD biomarkers. Taking these together, we have examined miRNAs participation in AD development, in particular the Pin1-targeted miRNAs.

We initially extracted candidate miRNAs from two most-cited online databases and reached to 3 candidates: miR296-5p, miR200c, miR200b. Every single candidate miRNA has been searched based on nine keywords in PubMed database. We successfully found miR-296-5p, mR874, miR200c, miR200b and miR140-5p as Pin1 targeting miRNAs, which negatively control its expression.

Considering that none of the profiles were from brain tissues, more convincing datasets is required to provide vigorous evidences, while there is only one such integrated dataset on GEO. Furthermore, it is important to examine differentially expressed candidate miRNAs pattern between AD and normal tissues to clearly demonstrate the miRNAs mediatory role in AD. We analyzed six miRNA expression profiles from several parts of the brain, CSF, and peripheral blood, among which the results are promising only for miR296-5p, that is being up-regulated in AD subjects.

Taking these together, we have identified some miRNAs mediating AD development through Pin1 control. Although our findings have some drawbacks considering the lack of sufficient and convincing proof as well as to-the-point studies, we believe our work would open new windows toward AD understanding and treatment.

Apparently, more experiments such as luciferase reporter assays and inhibition/transfection studies are necessary to promise the findings in the experimental stage. It is notable that we are going to run an experiment inhibiting 3 candidate miRNAs (miR296-5P, miR200c, and miR200b) employing antagomirs, in AD cell models to study possible therapeutic outcomes.

## VI. ACKNOWLEDGMENT

We appreciate Ms. T. Saraie’s kind helps in coordinating the team and Mr. S. Ansarin for providing the illustration of our pipeline. We express our deep gratitude to prof. De Stooper and Dr. P. Lau, and Dr. L. Cheng for sharing their data. Finally, we gratefully acknowledge Dr. Sharif Moradi for his valuable suggestions and discussions in our miRNA research. The authors declares no competing financial interests.

